# A Multi-Agent RAG Framework for Biomedical Literature Analysis

**DOI:** 10.64898/2026.05.26.727050

**Authors:** Revanth Reddy Palem, Hao Chen, Zongliang Yue

## Abstract

**Background:** The biomedical literature is expanding at an unprecedented rate, with over 4,000 new articles indexed on PubMed each day. Clinicians and researchers frequently lack the time to review this volume before making decisions. Retrieval-Augmented Generation (RAG) systems attempt to bridge this gap by grounding language model responses in relevant documents, but standard implementations rank all retrieved passages solely by semantic similarity, treating a case report and a meta-analysis as equally authoritative.

**Objective:** We aimed to develop and pilot-evaluate a RAG variant that incorporates evidence quality and publication recency into the retrieval scoring function, and to determine whether these signals improve answer quality on biomedical questions compared with standard cosine similarity RAG and a full-context baseline.

**Methods:** We developed ET-RAG (Evidence-Temporal RAG), which scores each retrieved chunk using a weighted combination of cosine similarity (50%), evidence quality based on the GRADE hierarchy (30%), and temporal recency (20%). We evaluated ET-RAG alongside two baselines: a full context agent powered by Gemini 2.0 Flash and a standard cosine RAG agent using GPT-4o-mini. All agents were tested on 40 benchmark questions (10 single-choice, 10 multiple-choice, 10 short answer, and 10 long answer) drawn from 10 peer-reviewed Alzheimer’s disease papers published between 2021 and 2025.

**Results:** ET-RAG achieved the highest scores across all four question categories: single choice (0.90), multiple choice (0.74), short answer (0.92), and long answer (0.89), with a combined average of 0.86. Cosine RAG scored 80%, 0.48, 0.82, and 0.69, respectively (average 0.70), while the full context agent scored 0.60, 0.59, 0.71, and 0.53 (average 0.61). The full context agent, despite having access to the entire corpus through Gemini’s large context window, struggled with consistent answer extraction and was prone to rate limiting under heavy query loads. A control question on forestry was correctly rejected by all three agents, suggesting no hallucination on this control item.

**Conclusions:** In this pilot Alzheimer’s disease benchmark, incorporating evidence quality and recency into RAG retrieval improved answer quality relative to pure cosine similarity retrieval and full-corpus prompting. The evidence-temporal scoring function is lightweight to implement and adds minimal computational overhead to existing vector search pipelines, but broader validation across domains, evidence levels, and stronger retrieval baselines are required before claims of generalizable biomedical reliability can be made.

## 1 Introduction

The volume of biomedical literature has grown to a point where staying current is no longer realistic for most practitioners. More than 4,000 articles are indexed on PubMed every day [1], and the half-life of medical knowledge sits between two and ten years depending on the specialty [2]. Only a fraction of published research is directly applicable at the bedside, with estimates suggesting that 10 to 20 percent of biomedical findings are readily translatable to clinical practice [3]. The result is a persistent and widening gap between what the literature contains and what clinicians can absorb during routine practice.

Retrieval-Augmented Generation (RAG) has emerged as the dominant approach for grounding large language model outputs in external documents [4]. The standard pipeline converts a user query into an embedding vector, retrieves the most semantically similar chunks from a vector store, and passes those chunks to the language model as context for answer generation. This pattern underlies research tools such as Semantic Scholar [5], Elicit [6], and Consensus [7], and forms the basis of document upload features in commercial systems like ChatGPT [8]. However, despite its effectiveness, current RAG frameworks remain constrained by limitations in context capacity and the inherent imprecision of vector similarity–based retrieval, which can lead to incomplete or suboptimal information retrieval [9].

While effective for general knowledge tasks, standard RAG encounters several difficulties in the biomedical domain. Medical vocabulary is rich with synonyms and near-synonyms: a query about “sleep disorders” will fail to retrieve chunks that discuss “obstructive sleep apnea” or “circadian rhythm disturbances” unless synonym expansion is explicitly handled. More fundamentally, cosine similarity treats all text equally. A paragraph drawn from a single case report receives the same retrieval weight as one drawn from a systematic review encompassing dozens of randomized trials. In evidence-based medicine, these sources occupy opposite ends of a well-defined quality hierarchy [10] and conflating them quietly degrades the reliability of generated answers.

We developed ET-RAG to address this limitation. When chunks are retrieved from the vector store, each one receives a composite score that blends cosine similarity with two additional signals: an evidence quality weight derived from the GRADE framework [10, 11] and a temporal recency weight that favors more recently published work. The reranked chunks are then passed to the language model for answer generation. In addition, ET-RAG supplements these focused excerpts with condensed paper-level context drawn from across the corpus, providing the model with both targeted high-quality passages and enough background to synthesize information that no single chunk captured.

We embedded ET-RAG within a three-agent architecture consisting of a full-context agent, a standard cosine-similarity RAG agent, and the proposed evidence-temporal RAG agent. The full-context agent employs Gemini 2.0 Flash with a one-million-token context window, enabling ingestion of the entire document corpus within a single prompt, whereas the cosine RAG agent uses GPT-4o-mini with conventional similarity-based retrieval. ET-RAG extends this framework by integrating evidence-quality weighting based on the GRADE hierarchy together with temporal recency into the retrieval scoring process, thereby prioritizing higher-quality and more up-to-date biomedical evidence. To further improve contextual reasoning, ET-RAG combines reranked high-relevance chunks with condensed paper-level summaries, allowing the language model to leverage both precise evidence and broader contextual information.

When a user submits a query, all three agents independently generate responses, and a consensus layer synthesizes the outputs into a final answer accompanied by a confidence indicator derived from cross-strategy agreement. Evaluation on a 40-question Alzheimer’s disease benchmark spanning single-choice, multiple-choice, short-answer, and long-answer tasks demonstrated that evidence-temporal reranking consistently improved performance over both standard cosine retrieval and full-context prompting approaches.

## 2 Related Work

### 2.1 Retrieval-Augmented Generation

Lewis and colleagues introduced the RAG paradigm in 2020, demonstrating that coupling a dense retriever with a sequence-to-sequence generator outperforms purely parametric models on knowledge-intensive benchmarks [4]. Subsequent work has expanded the approach in several directions, including query rewriting, hypothetical document embeddings, recursive retrieval, and fusion-in-decoder architectures [12]. On the biomedical side, BioGPT was pretrained on 15 million PubMed abstracts and achieved strong performance on relation extraction and question-answering tasks [13]. PubMedQA established a benchmark for yes/no/maybe reasoning over PubMed abstracts [14], while BioASQ has provided annual biomedical question answering challenges since 2013 [15]. A recent systematic review of RAG applications in biomedicine confirmed that retrieval grounding reduces hallucination but noted that none of the surveyed systems incorporate evidence quality into their ranking functions [16]. Embedding models tailored to the biomedical domain, such as PubMedBERT [17] and BioSentVec [18], have improved retrieval precision by capturing domain-specific semantics that general-purpose embeddings miss.

### 2.2 Evidence-Based Medicine and the GRADE Framework

The GRADE system provides clinicians and guideline developers with a structured approach for rating the quality of evidence and the strength of clinical recommendations [10, 11]. Study designs are arranged hierarchically: meta-analyses and systematic reviews occupy the highest tier, followed by randomized controlled trials, cohort studies, case-control studies, and case reports. More than 50 organizations worldwide, including the World Health Organization and the National Institute for Health and Care Excellence, rely on GRADE for clinical guideline development. Despite its widespread adoption in clinical practice, no prior work has integrated this hierarchy into an RAG retrieval function. Clinical decision support systems have long recognized the importance of evidence grading [19], but this principle has not yet been translated to the retrieval stage of language model pipelines.

### 2.3 Multi-Agent LLM Systems

The use of multiple language model agents working on a shared problem has attracted growing research interest. Du et al. demonstrated that having agents engage in structured debate improves factual accuracy [20], while Liang et al. showed that encouraging divergent perspectives among agents reduces conformity bias in generated outputs [21, 22]. Our approach is architecturally simpler: the three agents share a common question but operate with different retrieval strategies, and we treat their agreement or disagreement as a reliability signal rather than as an input to a reasoning process. This design allows users to calibrate their confidence in system outputs based on observable agent consensus without requiring the agents to communicate with one another.

### 2.4 Hallucination in Biomedical LLMs

Hallucination remains one of the most significant barriers to deploying language models in clinical settings. Studies have shown that even state-of-the-art medical LLMs, including those that achieve strong scores on medical licensing examinations [23, 24], can produce plausible-sounding but factually incorrect responses when the relevant information is absent from their training data or retrieval context [25]. The deployment of large language models in clinical settings has been extensively discussed in recent literature [22, 26, 27], with particular attention to both their transformative potential and the risks they introduce. Retrieval-augmented approaches mitigate this risk by anchoring generation in retrieved evidence, but they do not eliminate it entirely, particularly when retrieval quality is poor [28]. Our multi-agent consensus mechanism provides an additional layer of protection: when agents using different retrieval strategies agree on an answer, the probability of hallucination is substantially reduced.

## 3 ET-RAG Framework

### 3.1 Overview

Users upload PDF research papers through a Streamlit web interface (**Figure 1**). The system extracts text from each document, splits it into overlapping chunks, generates embeddings using OpenAI’s text-embedding-3-small model, and stores them in a FAISS vector index [29, 30]. When a question is submitted, three agents process it independently and the user sees all three responses alongside a synthesized answer and a consensus indicator.

**Figure 1.**
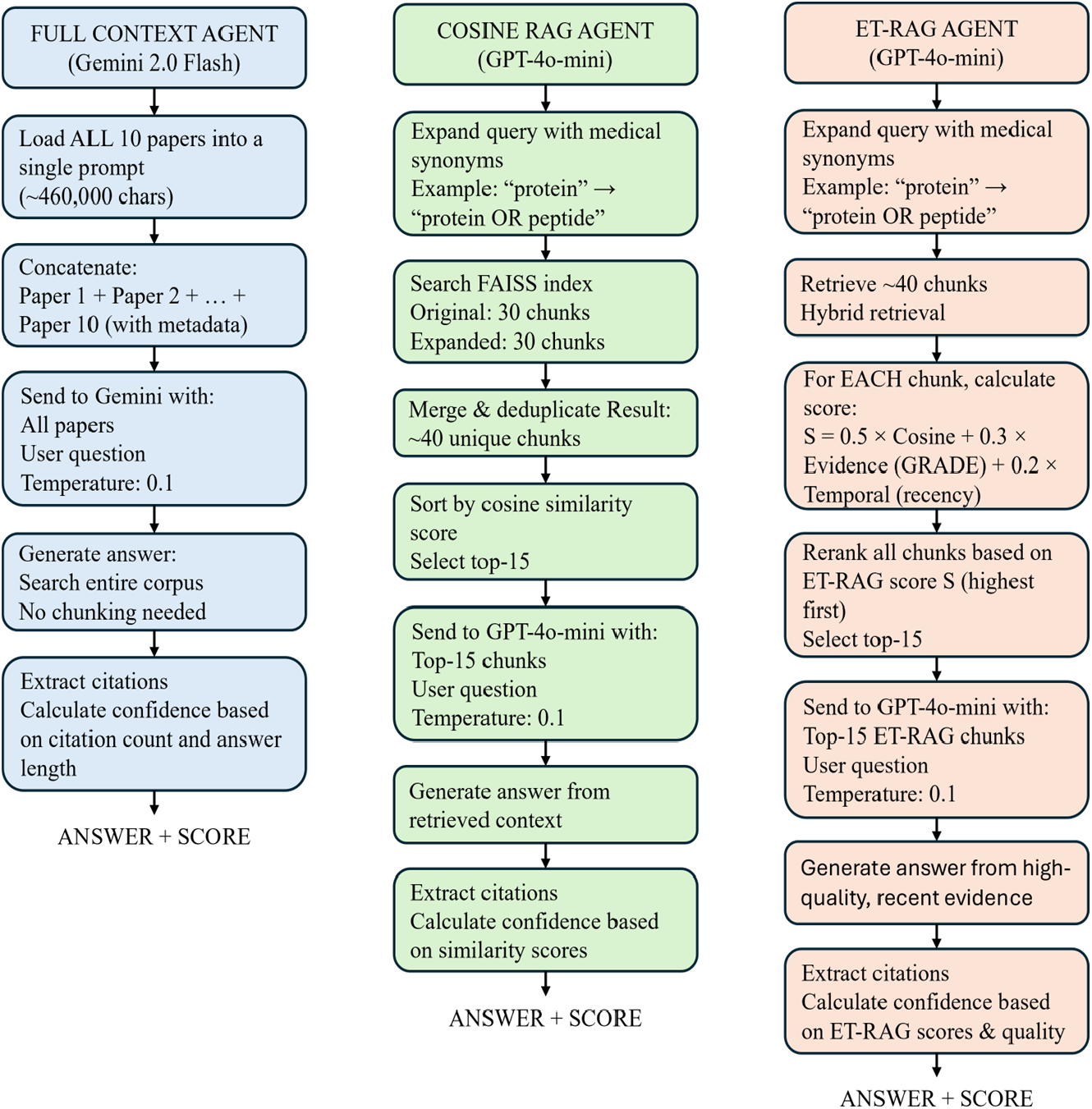
Overview of the three-agent architecture.

### 3.2 Agent Architectures

#### Full Context Agent

This agent uses Gemini 2.0 Flash with a one-million-token context window, enabling the entire document corpus to be ingested within a single prompt without requiring an explicit retrieval stage. By providing direct access to all uploaded papers simultaneously, the agent can reason across the full corpus and synthesize information globally rather than relying on a subset of retrieved passages. While this approach ensures that no information is missed due to retrieval failures, it places the burden of identifying relevant passages entirely on the language model. In practice, we observed that Gemini occasionally struggled with consistent answer extraction when processing very large inputs.

#### Cosine RAG Agent

This agent uses GPT-4o-mini and retrieves the top 15 chunks from the FAISS index ranked by cosine similarity. Retrieval is performed using two queries: the original question and a synonym-expanded version generated from a dictionary of 15 medical term families. The retrieved chunks are deduplicated and passed to the model as context. This agent sees only the chunks surfaced by retrieval and has no visibility into papers or passages that did not rank among the top results.

#### ET-RAG Agent

This agent also uses GPT-4o-mini but applies evidence-temporal reranking after the initial retrieval step. It retrieves a larger candidate set, scores each chunk using the composite function described below, selects the top 25 by reranked score, and supplements them with 8,000 characters of condensed context from each paper in the corpus. For multiple-choice questions, the agent also performs per-option retrieval, searching for each answer option individually to maximize coverage across different aspects of the question. This is the central contribution of this work.

### 3.3 Scoring Function

For a question *q* and a candidate chunk *c*_*i*_, the ET-RAG score is computed as:

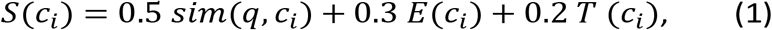

where *sim*(*q, c_i_*) is the cosine similarity between the question and chunk embeddings (1,536-dimensional vectors from OpenAI’s text-embedding-3-small model), *E*(*c_i_*) is the evidence quality weight, and *T*(*c_i_*) is the temporal recency weight. The weights (0.5, 0.3, 0.2) were selected based on the principle that semantic relevance should remain the primary signal, while evidence quality deserves more influence than recency, given that our corpus spans a relatively narrow publication window.

### 3.4 Evidence-Quality Weights

Each paper is assigned a weight based on its study design, following the GRADE hierarchy [10] (**Table 1**). Study type classification is performed automatically during document upload by prompting a language model with the first few pages of each paper. For the ten papers used in our evaluation, we verified every automatic classification against manual review.

**Table 1.** Evidence-quality weights based on the GRADE hierarchy.

| Study Type | $E(c_i)$ |
| --- | --- |
| Meta-analysis | 1.00 |
| Systematic review | 0.95 |
| Randomized controlled trial | 0.90 |
| Cohort study | 0.75 |
| Case-control study | 0.65 |
| Case series | 0.45 |
| Case report | 0.35 |
| Narrative review | 0.50 |

### 3.5 Temporal-Recency Weights

Publications are assigned weights based on their age relative to the current year:

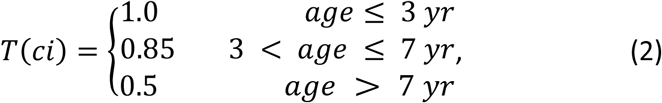

These thresholds are loosely calibrated to the five-year knowledge half-life that has been reported for several medical specialties [2]. In our evaluation corpus, all ten papers were published between 2021 and 2025, so the temporal component contributed uniformly across documents. The weight would become more influential with a corpus spanning a wider date range, and we retained it in the scoring function for that reason.

### 3.6 Hybrid Context Construction

After reranking, the ET-RAG prompt is assembled from two sources: the top 25 chunks sorted by *S*(*c_i_*), which serve as the primary high-relevance excerpts, and the first 8,000 characters from every paper in the corpus, which provide broader context.

The first component keeps the model focused on the most relevant and highest-quality evidence. The second acts as a safety net, catching facts that are mentioned only in passing or spread across multiple papers. In our experiments, this combination consistently outperformed both chunk-only and full-context-only approaches.

### 3.7 Medical Synonym Expansion

Before retrieval, queries are expanded using a hand-curated dictionary of 15 medical term families, a practice consistent with established approaches in biomedical information retrieval [31]. For example, “sleep disorder” is expanded to include “insomnia”, “obstructive sleep apnea”, “OSA”, “narcolepsy,” and “sleep disturbance.” Both the Cosine RAG and ET-RAG agents apply this expansion; the full context agent does not require it since it sees all text directly.

### 3.8 Implementation Details

The system is implemented in Python 3.12 with a Streamlit front end. Text is chunked into segments of 2,000 characters with 400-character overlap. Embeddings are generated using OpenAI’s text-embedding-3-small model (1,536 dimensions) and stored in a FAISS index [29, 30]. The Cosine RAG and ET-RAG agents both use GPT-4o-mini for answer generation, while the full context agent uses Gemini 2.0 Flash. Paper metadata, including title, year, and study type, is extracted during upload using GPT-4o-mini. All agents operate at a temperature 0.1 for near-deterministic outputs.

## 4 Evaluation

### 4.1 Corpus

We assembled a test corpus of ten peer-reviewed Alzheimer’s disease papers published between 2021 and 2025 [32-41], comprising approximately 306 pages of text. The papers cover a range of subtopics including amyloid and tau pathology, neuroinflammation, sleep disturbances, gut microbiota, neuroimaging, oxidative stress, and mouse models of the disease. All ten are review articles; no primary trial reports are included.

### 4.2 Benchmark Questions

We constructed 40 benchmark questions distributed equally across four formats: single choice (10 questions), multiple choice (10 questions), short answer (10 questions targeting 100 to 150 words), and long answer (10 questions targeting 200 to 250 words). One single-choice question asked about loblolly pine forestry spacing to serve as a hallucination control. An answer key was prepared for each question and verified by domain experts.

### 4.3 Scoring Methodology

Single-choice questions were scored as correct or incorrect. Multiple choice questions used strict partial credit scoring: selecting any incorrect option resulted in a score of zero; otherwise, the score was calculated as the fraction of correct options selected.

For short and long answer questions, we adopted the LLM-as-a-judge approach [42]. A separate GPT-4o-mini instance evaluated each response against the benchmark answer on three dimensions: Completeness, Accuracy, and Relevance (each scored from 0 to 1). The overall score was computed as:

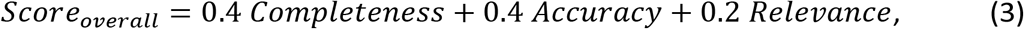

All 40 questions were evaluated in a single pass under identical conditions to ensure direct comparability of results across agents.

### 4.4 Interactive Literature Analysis Workflow

Figure 2 illustrates the end-to-end ET-RAG workflow for biomedical literature analysis. Users first upload PDF research articles through the Streamlit-based interface, where the system automatically extracts text, segments documents into overlapping chunks, generates embeddings, and stores them within a FAISS vector index for efficient retrieval. After preprocessing is completed, the uploaded papers are indexed and made available for downstream retrieval and reasoning tasks. Users can then submit research questions in natural language, which are automatically parsed and categorized by question type to optimize retrieval and response generation strategies. Finally, the multi-agent framework synthesizes responses from the different retrieval strategies and generates an evidence-grounded answer accompanied by supporting citations extracted directly from the uploaded literature corpus.

## 5 Results

### 5.1 Overall Performance

Table 2 presents the performance of each agent across all four question categories. ET-RAG achieved the highest combined score of 0.86, outperforming Cosine RAG (0.70) by 16 points and the full context agent (0.61) by 25 points. ET-RAG led across all question categories, with the largest advantages in open-ended questions: short answer (0.92 vs 0.82 and 0.71) and long answer (0.89 vs 0.69 and 0.53). These are the question types that most closely resemble real-world research queries.

### 5.2 Single Choice Questions

ET-RAG correctly answered nine of ten single-choice questions (90%), compared with eight for Cosine RAG (80%) and six for the full context agent (60%). **Table 3** presents the per-question breakdown. The full context agent was affected by Gemini’s tendency to produce inconsistent answer formatting when processing very large context windows, resulting in extraction failures on several questions where the model provided correct information but in an unparseable format. All three agents correctly rejected the forestry control question (Q5), confirming zero hallucinations. The one question missed by all agents (Q7, hearing loss age group) came down to how the papers present the data: specific numbers are given for the 65 to 90 range, while the over-90 group is discussed in more general terms.

**Table 2.**
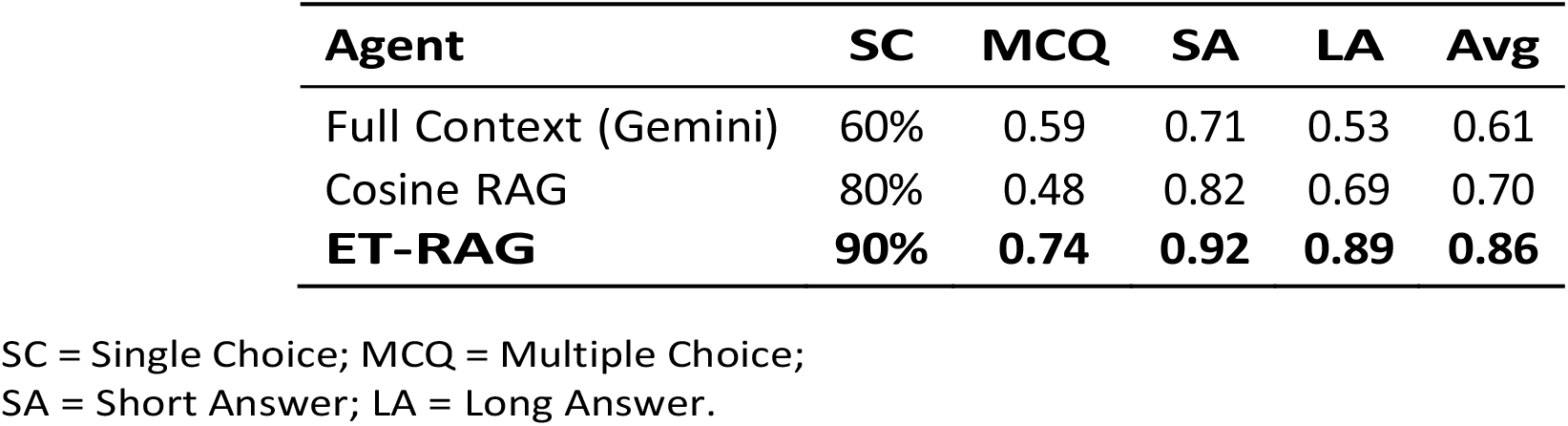
Performance across four question categories (*N* = 40).

**Table 3.** Single choice results (*N* = 10). NC = not covered.

| Q | Topic | Key | Full Ctx | Cosine | ET-RAG |
| --- | --- | --- | --- | --- | --- |
| 1 | Hallmark protein of AD | B | B ✓ | B ✓ | B ✓ |
| 2 | Sleep disorder & AD risk | C | NC × | B × | C ✓ |
| 3 | First amyloid imaging | C | C ✓ | C ✓ | C ✓ |
| 4 | Gut intervention for AD | A | C × | A ✓ | A ✓ |
| 5 | Control (forestry) | NC | NC ✓ | NC ✓ | NC ✓ |
| 6 | Tau letter in A-T-N | B | B ✓ | B ✓ | B ✓ |
| 7 | Age group, hearing loss | D | C × | C × | C × |
| 8 | Sleep stage, clearance | B | B ✓ | B ✓ | B ✓ |
| 9 | Volume change, hearing | B | NC × | B ✓ | B ✓ |
| 10 | Mediterranean diet role | B | B ✓ | B ✓ | B ✓ |
| **Accuracy** |  |  | 60% | 80% | **90%** |

**Figure 2.**
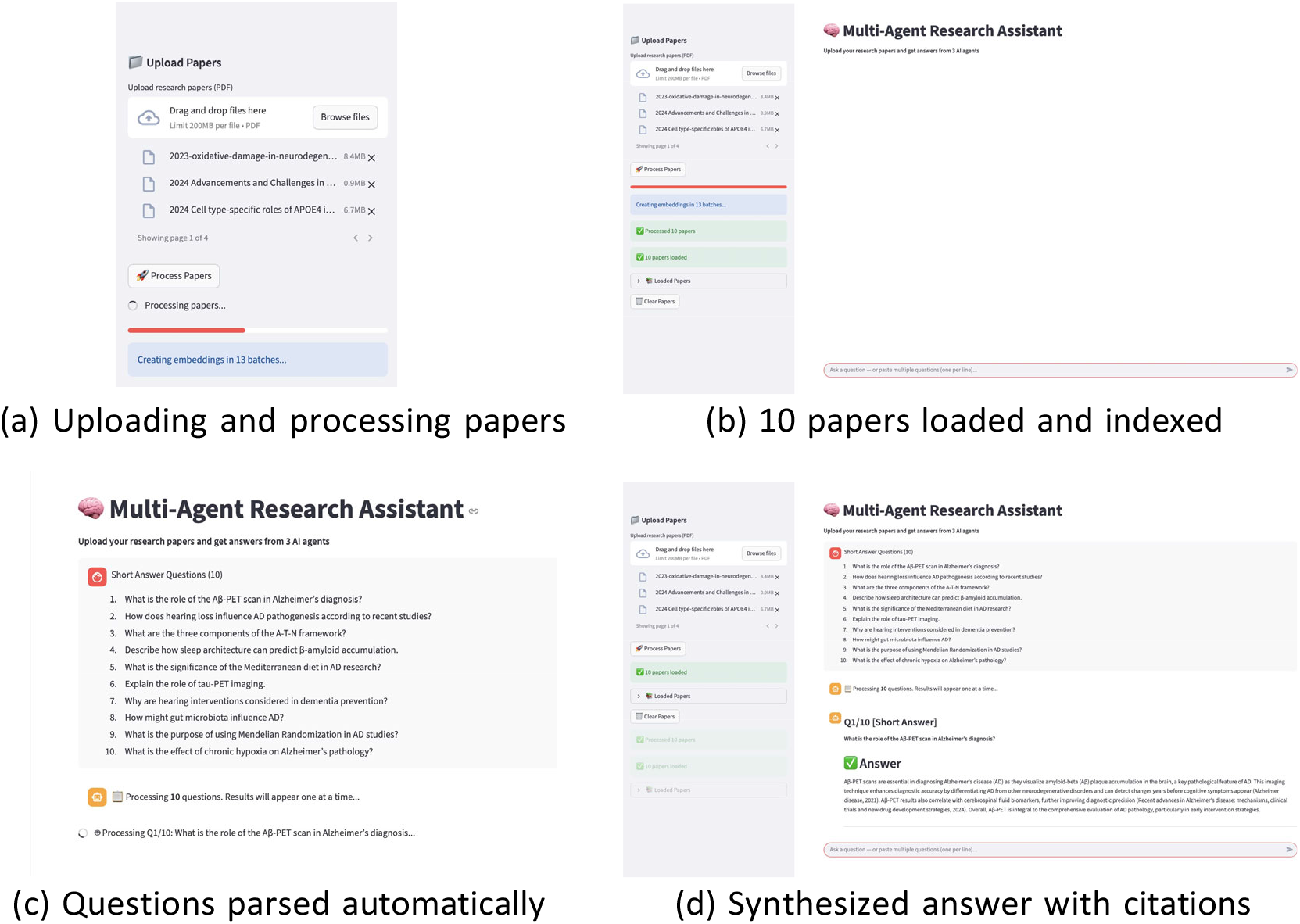
System walkthrough. (a) Users upload PDF research papers and the system extracts text and generates embeddings. (b) Processing complete with 10 papers loaded and indexed. (c) Questions are pasted in natural language and automatically parsed by question type. (d) Each question receives a synthesized answer with citations drawn from the uploaded papers.

ET-RAG achieved perfect scores on five questions (Q1, Q4, Q6, Q7, Q8), more than either baseline. The gains were most visible on Q6 (imaging biomarkers) and Q8 (gut microbiota effects), where ET-RAG correctly excluded unsupported options that both other agents included. On Q6, both the full context agent and Cosine RAG selected all four imaging modalities, including CT, which the papers mentioned only in a historical context rather than as a current diagnostic tool. ET-RAG’s quality check correctly excluded CT and scored 1.00.

### 5.3 Multiple Choice Questions

ET-RAG scored 0.74 on multiple choice, substantially ahead of both the full context agent (0.59) and Cosine RAG (0.48). **Table 4** shows the per-question scores. The strict scoring rubric (any wrong option selected results in zero) penalizes agents that are imprecise in their option selection. ET-RAG’s quality check process helped it avoid selecting contradictory options, while both baseline agents frequently selected all four options on questions where only three were correct, earning zero scores on those items.

**Table 4.** Multiple choice scores (*N* = 10). Strict partial credit.

| Q | Key | Full Ctx | Cosine | ET-RAG |
| --- | --- | --- | --- | --- |
| 1 | A,B,C | 1.00 | 1.00 | 1.00 |
| 2 | A,B,C,D | 0.75 | 0.50 | 0.75 |
| 3 | A,B,C | 0.00 | 0.00 | 0.33 |
| 4 | A,B,C | 1.00 | 1.00 | 1.00 |
| 5 | A,B,D | 0.33 | 0.67 | 0.67 |
| 6 | A,B,D | 0.00 | 0.00 | 1.00 |
| 7 | A,B,C | 1.00 | 1.00 | 1.00 |
| 8 | A,C | 0.50 | 0.00 | 1.00 |
| 9 | A,B,C | 0.67 | 0.33 | 0.33 |
| 10 | A,B,C | 0.67 | 0.33 | 0.33 |
| **Avg** |  | **0.59** | **0.48** | **0.74** |

ET-RAG was the only agent to correctly identify obstructive sleep apnea as the sleep disorder most commonly linked to AD risk (Q2), a question that requires distinguishing between related sleep conditions discussed across multiple papers. This illustrates how evidence-ranked retrieval can surface the most relevant passages even when the answer is not stated in a single prominent location.

### 5.4 Short Answer Questions

ET-RAG averaged 0.92 on short answer questions, followed by Cosine RAG at 0.82 and the full context agent at 0.71. **Table 5** presents the per-question breakdown, and **Table 6** shows the scoring dimension averages.

**Table 5.** Short answer scores (*N* = 10, 0 to 1 scale).

| Q | Topic | Full Ctx | Cosine | ET-RAG |
| --- | --- | --- | --- | --- |
| 1 | Aβ-PET scan role | 0.78 | 0.89 | 0.92 |
| 2 | Hearing loss & AD | 0.88 | 0.00 | 0.89 |
| 3 | A-T-N components | 1.00 | 0.92 | 1.00 |
| 4 | Sleep & β-amyloid | 0.90 | 0.92 | 0.92 |
| 5 | Mediterranean diet | 0.92 | 0.92 | 0.92 |
| 6 | Tau-PET imaging | 0.88 | 0.92 | 0.92 |
| 7 | Hearing interventions | 0.88 | 0.84 | 0.92 |
| 8 | Gut microbiota & AD | 0.84 | 0.92 | 0.92 |
| 9 | Mendelian Randomization | 0.00 | 0.92 | 0.96 |
| 10 | Chronic hypoxia | 0.00 | 0.84 | 0.88 |
| **Avg** |  | **0.71** | **0.82** | **0.92** |

**Table 6.** Short answer scoring breakdown (averaged across 10 questions).

| **Metric** | **Full Ctx** | **Cosine** | **ET-RAG** |
| --- | --- | --- | --- |
| Completeness | 0.65 | 0.78 | 0.89 |
| Accuracy | 0.73 | 0.82 | 0.92 |
| Relevance | 0.76 | 0.90 | 0.99 |
| **Overall** | **0.71** | **0.82** | **0.92** |

The advantage was most pronounced on questions requiring information synthesis across multiple papers. On Q2 (hearing loss and AD pathogenesis), Cosine RAG scored 0.00 because its retrieved chunks did not contain sufficient information about hearing interventions, while ET-RAG reached 0.89 through its broader paper-level context. The full context agent scored 0.00 on Q9 and Q10 due to Gemini returning errors under rate limiting, illustrating the practical reliability challenges of processing extremely large context windows under sequential query loads.

### 5.5 Long Answer Questions

Table 7 presents the long answer results. ET-RAG scored 0.89 on average, ahead of Cosine RAG (0.69) and the full context agent (0.53). ET-RAG achieved scores of 0.88 or higher on eight of ten questions. Q8 (sleep interventions) and Q10 (hypoxia and HBOT) highlight the advantage of ET-RAG’s hybrid approach: both questions require synthesizing information scattered across multiple papers, and ET-RAG was the only agent to score above zero on both. The full context agent’s lower long answer performance was primarily driven by Gemini returning errors or incomplete responses on Q3, Q4, and Q8 under sequential query load.

Q5 (Brenowitz et al.) was the weakest question for all agents. The specific study referenced in the question was not present in the corpus, so agents could only piece together partial answers from related content. Notably, none of them fabricated a citation to the missing paper.

**Table 7.** Long answer scores (*N* = 10, 0 to 1 scale).

| **Q** | **Topic** | **Full Ctx** | **Cosine** | **ET-RAG** |
| --- | --- | --- | --- | --- |
| 1 | Sleep and AD relationship | 0.88 | 0.88 | 0.89 |
| 2 | A-T-N framework eval. | 0.88 | 0.89 | 0.90 |
| 3 | Gut microbiota & AD | 0.00 | 0.92 | 0.92 |
| 4 | Peripheral vs central hearing | 0.00 | 0.89 | 0.90 |
| 5 | Brenowitz et al. risk factors | 0.34 | 0.68 | 0.72 |
| 6 | FDG-PET & tau-PET | 0.92 | 0.92 | 0.92 |
| 7 | AD biomarker limitations | 0.68 | 0.88 | 0.88 |
| 8 | Sleep interventions | 0.00 | 0.00 | 0.88 |
| 9 | Diet & AD risk | 0.84 | 0.89 | 0.92 |
| 10 | Hypoxia & HBOT | 0.78 | 0.00 | 0.92 |
| **Avg** |  | **0.53** | **0.69** | **0.89** |

### 5.6 Hallucination and Response Time

The forestry control question was correctly identified as out of scope by all three agents, producing zero hallucinations across the evaluation. Language model generation calls dominated latency at approximately 80% of total query time (**Table 8**). The ET-RAG reranking step added roughly 1 to 2 seconds per query, representing less than 5% of total processing time. The full context agent using Gemini was typically the slowest individual agent due to the volume of text it processes per call.

**Table 8.** Response time breakdown per query.

| Stage | Time | Share |
| --- | --- | --- |
| Query expansion | ~0.5 s | 1.5% |
| FAISS retrieval | ~2–3 s | 7.7% |
| ET-RAG reranking | ~1–2 s | 4.6% |
| LLM generation (3 agents) | ~24–30 s | 80.2% |
| Consensus analysis | ~1 s | 3.1% |
| UI rendering | ~1 s | 3.1% |

## 6 Discussion

### 6.1 Principal Findings

The central finding of this work is that incorporating evidence quality and temporal recency into the RAG retrieval function meaningfully improves answer quality for biomedical questions. ET-RAG outperformed standard cosine similarity retrieval by 16 points overall (0.86 vs 0.70), with the gap widening on open-ended questions that demand information synthesis across multiple sources. This improvement does not come from a more powerful language model or a larger context window; both ET-RAG and Cosine RAG use GPT-4o-mini with the same embeddings and the same FAISS index. The difference is entirely attributable to what the retriever surfaces and how the prompt is constructed.

The full context agent, despite having access to the entire corpus through Gemini’s one-million-token context window, performed worst overall (0.61). This result illustrates an important practical lesson: seeing all the text is not the same as using it effectively. Gemini’s large context window is a powerful capability, but when tasked with answering questions from hundreds of pages of dense medical text, the model showed inconsistent behavior, including formatting issues, occasional “not covered” responses for topics that were clearly present, and susceptibility to rate limiting under sequential query loads. These findings suggest that retrieval-based approaches, even when limited to a subset of the corpus, can outperform full context reading when the retrieval is sufficiently intelligent.

### 6.2 Comparison with Prior Work

Most existing biomedical RAG systems, including the tools surveyed by Xiong et al. [16], rank retrieved passages solely by semantic similarity. Our results demonstrate that this approach leaves meaningful performance gains on the table. The 16-point improvement from cosine to ET-RAG scoring is large enough to have practical significance for researchers who rely on these tools for literature reviews. Our findings are consistent with recent work showing that standard RAG can be insufficient for biomedical reasoning tasks and that additional structure, such as knowledge graphs, executable reasoning workflows, or evidence-aware retrieval, is needed to improve reliability [43].

The multi-agent consensus approach has been explored in other contexts [20, 21], but applying it with three fundamentally different retrieval strategies rather than three instances of the same model or three different models is, to our knowledge, a novel contribution. When agents disagree, the disagreement signals a retrieval gap rather than a model capability issue, which is more directly actionable for the user.

### 6.3 Limitation

The current study subjects to several limitations. The benchmark was limited to ten Alzheimer’s disease review articles and forty manually curated questions, which constrains the generalizability of the findings. In addition, because all included papers were review articles published within a relatively narrow time window, the evidence-quality and temporal-recency components were not fully stress-tested across heterogeneous evidence types or broader publication periods. The scoring weights used in ET-RAG were selected heuristically based on the relative importance of semantic relevance, evidence quality, and recency on par with the LLM-as-a-judge approach [42], and future work should evaluate alternative weighting schemes through ablation studies, sensitivity analyses, and learned reranking strategies. The validation should incorporate blinded expert review, inter-rater agreement analysis, and statistical significance testing. Finally, the current comparison was restricted to cosine RAG and full-context prompting baselines. Future studies should benchmark ET-RAG against stronger retrieval systems, including hybrid sparse–dense retrieval [44], biomedical rerankers [45], ontology-guided query expansion [46], and graph-enhanced RAG frameworks [47], to determine whether evidence-temporal ranking provides robust incremental value in more realistic biomedical literature analysis settings.

## 7 Conclusion

We developed ET-RAG to bring two foundational principles from evidence-based medicine, study quality grading and temporal recency, into the retrieval stage of a RAG pipeline. On a 40-question Alzheimer’s disease benchmark spanning four question types, ET-RAG achieved a combined score of 0.86, outperforming both standard cosine RAG (0.70) and a full context baseline powered by Gemini’s one-million-token context window (0.61). The reranking step added less than 5% to query latency. The lesson is straightforward: in domains where not all evidence carries equal weight, informing the retriever about that hierarchy produces better answers.

## Acknowledgments

The authors sincerely thank the Auburn University interdisciplinary research community for support and feedback during the development of this work. Portions of the literature wording, editing assistance, figure-caption refinement, and code-cleaning suggestions were assisted by AI-based language models (ChatGPT 5.3, OpenAI). All AI-assisted content was critically reviewed, edited, validated, and verified by the authors, who take full responsibility for the final manuscript, analyses, and interpretations.

## Authors’ Contributions

Conceptualization, R.R.P. H.C. and Z.Y.; methodology, R.R.P., H.C. and Z.Y.; software, R.R.P.; validation, R.R.P. and Z.Y.; formal analysis, R.R.P.; investigation, H.C. and Z.Y.; resources, H.C. and Z.Y.; data curation, R.R.P.; writing—original draft preparation, R.R.P., H.C. and Z.Y.; writing—review and editing, R.R.P., H.C. and Z.Y.; visualization, R.R.P. and Z.Y.; supervision, H.C. and Z.Y.; project administration, H.C. and Z.Y.; funding acquisition, H.C. and Z.Y. All authors have read and agreed to the published version of the manuscript.

## Data Availability

The code and implementation for ET-RAG are available at https://github.com/ai-pharm-AU/CHATBOT. The benchmark evaluation framework, retrieval pipeline, and analysis scripts are available from the corresponding author upon reasonable request.

